# A new method for inferring timetrees from temporally sampled molecular sequences

**DOI:** 10.1101/620187

**Authors:** Sayaka Miura, Koichiro Tamura, Sergei L. Kosakovsky Pond, Louise A. Huuki, Jessica Priest, Jiamin Deng, Sudhir Kumar

## Abstract

Pathogen timetrees are phylogenies scaled to time. They reveal the temporal history of a pathogen spread through the populations as captured in the evolutionary history of strains. These timetrees are inferred by using molecular sequences of pathogenic strains sampled at different times. That is, temporally sampled sequences enable the inference of sequence divergence times. Here, we present a new approach (RelTime with Dated Tips [RTDT]) to estimating pathogen timetrees based on the relative rate framework underlying the RelTime approach. RTDT does not require many of the priors demanded by Bayesian approaches, and it has light computing requirements. We found RTDT to be accurate on simulated datasets evolved under a variety of branch rates models. Interestingly, we found two non-Bayesian methods (RTDT and Least Squares Dating [LSD]) to perform similar to or better than the Bayesian approaches available in BEAST and MCMCTree programs. RTDT method was found to generally outperform all other methods for phylogenies in with autocorrelated evolutionary rates. In analyses of empirical datasets, RTDT produced dates that were similar to those from Bayesian analyses. Speed and accuracy of the new method, as compared to the alternatives, makes it appealing for analyzing growing datasets of pathogenic strains. Cross-platform MEGA X software, freely available from http://www.megasoftware.net, now contains the new method for use through a friendly graphical user interface and in high-throughput settings.

**AUTHOR SUMMARY:** Pathogen timetrees trace the origins and evolutionary histories of strains in populations, hosts, and outbreaks. The tips of these molecular phylogenies often contain sampling time information because the sequences were generally obtained at different times during the disease outbreaks and propagation. We have developed a new method for inferring timetrees for phylogenies with tip dates, which improves on widely-used Bayesian methods (e.g., BEAST) in computational efficiency and does not require prior specification of population parameters, branch rate model, or clock model. We performed extensive computer simulation and found that RTDT performed better than the other methods for the estimation of divergence times at deep node in phylogenies where evolutionary rates were autocorrelated. The new method is available in the cross-platform MEGA software package that provides a graphical user interface, and allows use via a command line in scripting and high throughput analysis (www.megasoftware.net).

## Introduction

Molecular phylogenetics enables dating of the origin of pathogens and of the emergence of new strains [1–3]. Typically, strains are sampled from individuals and populations during an ongoing or historical outbreak [4–9]. When sequences are paired with their sampling times, it becomes possible to calibrate molecular phylogenies of pathogen sequences and infer the timing of pathogen evolution. For example, HIV-1 sequences have been sampled at various times and geographic locations following its initial characterization in 1983 [2, 9, 10]. Analyses of sequences extracted from circulating strains and “archived” strains from preserved tissue samples have established that HIV-1 (group M) entered the human populations in the early 20^th^ century in Sub-Saharan Africa [10] and that subsequently dispersed across the globe [11, 12].

Many competing methods are available to build pathogen timetrees that estimate the timing of divergence of lineages in the tree [13–21]. These methods start with the evolutionary tree of sequences and build timetrees using the information on sequence sampling times, provided that the tips in the phylogeny are not contemporaneous. In these analyses, sampling times serve as calibrations that provide a means to date historical sequence divergences. These analyses are different from those used for the estimation of species divergence times because the sampling times of sequences from different species are effectively simultaneous. The difference in the sampling years for all sequences in interspecies datasets can be assumed to be effectively zero when compared to the time-scale of speciation.

The Bayesian framework underlies many of the widely-used tools for building pathogen timetrees (MCMCTree [15] and BEAST [14]). The use of Bayesian methods requires researchers to specify a clock prior that governs the change of evolutionary rate over lineages and a coalescent model (demographic history or birth-and-death) to generate a tree prior and compute likelihoods [14, 15]. Such information is rarely available *a priori*, and time estimates can vary when using different priors [22], resulting in alternative biological interpretations [15, 23]. Also, evolutionary processes that are not adequately modeled in the standard frameworks. For example natural selection or severe heterotachy can severely distort rate estimates and produce inferences that are contradicted by historical records or other sources of calibration information, e.g. endogenous retroviral elements [24–26].

Here, we present an approach based on the relative rate framework underlying the RelTime method [27, 28]. The RelTime method is not computationally demanding and it does not require explicit clock and coalescent model priors. Both simulated and empirical analyses have shown RelTime to perform well for dating species evolution [27, 28]. The new approach advances RelTime by relaxing the requirement that all tips in the phylogenetic tree are contemporaneous (i.e., sampling time *t* = 0), making it suitable for dating of pathogenic strains. We call it the RelTime with Dated Tips (RTDT) approach. Similar to RelTime, RTDT does not require one to pre-specify rate models (e.g., autocorrelated vs. independent and exponential vs. lognormal) or a population dynamics model.

Through the analysis of simulated datasets generated under different assumptions and empirically derived phylogenies, we compared the accuracy of dates estimated by RTDT with Bayesian (BEAST [14] and MCMCTree [15]) and non-Bayesian (Least Squares Dating, LSD [16]) methods. We chose these three methods, because they have been used in sequence data analysis. In the past, some studies have reported the accuracy of estimation of substitution rates or the age of the root node of phylogeny [13, 20]. However, the accuracy of node-by-node age estimates remains to be evaluated. To et al. [16] conducted computer simulations, but only reported the average of the absolute difference in actual and estimated times for all the nodes in a phylogeny to compare methods. This measure does not detect node-specific biases and patterns. Also, previous computer simulation studies have only tested independent branch rate (IBR) model, so the performance is not known for phylogenies in which branch rates are autocorrelated (ABR model). This is important because the ABR model fits inter-species data sets much better [29] and may actually provide a better fit for the viral phylogenies as well. Therefore, much remains to be learned about the performance of molecular dates obtained by using previous Bayesian and non-Bayesian methods. Here, we present a new method and extensive computer simulation evaluation of Bayesian and non-Bayesian methods to yield new, unique insights into the performance of tip-dating methods in building pathogen timetrees.

## RESULTS

### New Approach (RTDT) for estimating divergence times using temporally sampled sequences

We illustrate the new approach by using a simple example dataset containing four ingroup sequences (*x*_1_, *x*_2_, *x*_3_, *x*_4_) with an outgoup sequence (Fig. 1A), where RTDT requires a phylogeny with outgroup specified. This is different from Bayesian methods (e.g., those implemented in BEAST), which jointly estimate phylogenies and divergence times without requiring the specification of outgroup sequences. In the ingroup, sequence *x*_*i*_ is assumed to be sampled in the year of *t*_*i*_ (2001, 2003, 2002, and 2011, for *x*_1_, *x*_2_, *x*_3_, and *x*_4_, respectively) and *b*_*i*_ are the branch lengths, expressed in expected substitutions per site (Fig. 1A). The goal is to estimate the time at internal nodes, X, Y, and XY: *t*_X_, *t*_Y_, and *t*_XY_.

**Figure 1.**
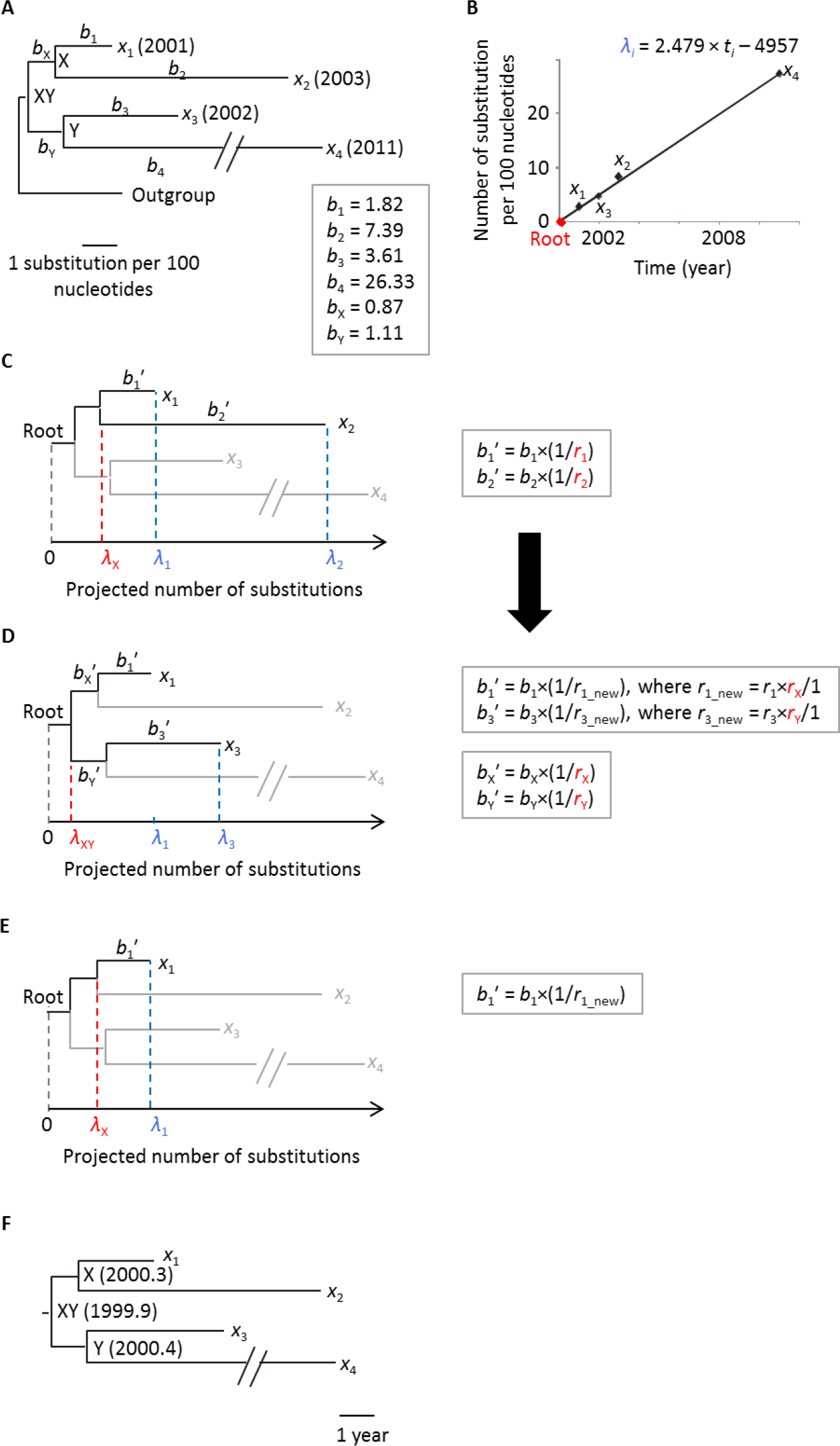
RelTime with Dated-Tips (RTDT) approach. (A) A phylogeny of five pathogen sequences (*x*_1_, *x*_2_, *x*_3_, *x*_4_, and outgroup), with branch lengths (*b*_*i*_). The year of sequence sampling (*t*_i_) is given in the parenthesis. The internal nodes are indicated by X, Y, and XY. (B) The relationship between the path lengths from node XY to tip and sampling times. For example, the point of *x*_1_ is (2001, *b*_X_ + *b*_1_). In the current example, the linear regression expression is *λ*_*i*_ = 2.479 × *t*_*i*_ − 4957. We locate a root at the position of *λ* = 0 along the regression line. (C-E) Projected phylogeny. A root-to-tip lengths were projected using linear regression. We first estimate relative rates at *b*_1_-*b*_4_, i.e., *r*_1_-*r*_4_ (C), and then estimate those at deeper positions of the phylogeny, i.e., *r*_X_ and *r*_Y_ (D). Lastly, we estimate the projected length from root to internal nodes, e.g., *λ*_X_ (E). (F) Estimated timetree. The final divergence times are estimated by using the regression line in panel B.

This phylogeny has a time-scale measured in chronological time (*t*_*i*_) and the number of substitutions (*b*_*i*_). In the RTDT approach, we first project the path length *λ*_*i*_ (number of substitutions) from the root to a tip (*x*_*i*_) of the phylogeny under the assumption that *x*_*i*_ accumulated substitutions to the year of the sampling time, *t*_*i*_, with a constant evolutionary rate (Fig. 1B). The projection is accomplished by first regressing the estimated length (in substitutions/site) from the node ingroup latest common ancestor (XY) to a tip (*x*_i_) in the original tree using the corresponding sampling time. This slope is used to project root-to-tip length, *λ*_*i*_, forward in time. In our example, *λ*_*i*_ = 2.479 × *t*_*i*_ − 4957. For example, the projected root-to-node length for sequence *x*_1_ is *λ*_1_ = 2.479 × 2001 − 4957 = 3.48. Note that the root in this projection is an “internal-root,” which is located at the position of zero substitution along the slope (Fig. 1B).

If the evolutionary rate were shared between branches *b*_1_ and *b*_2_, then the length from root to the internal node X, i.e., *λ*_X_, predicted by using *λ*_1_ and *b*_1_ and that predicted by using *λ*_2_ and *b*_2_ should be the same. In practice, they are not the same: *λ*_X_ is predicted to be 1.66 when using *λ*_1_ and *b*_1_ (= *λ*_1_ − *b*_1_ = 3.48 − 1.82) and 1.05 when using *λ*_2_ and *b*_2_ (= *λ*_2_ − *b*_2_ = 8.44 − 7.39), respectively. This suggests the inequality of evolutionary rates between *b*_1_ and *b*_2_. Under the RRF framework [27, 28] we, therefore, estimate their relative rates, *r*_1_ and *r*_2_, respectively, in which these two sister lineages inherited rates from their common ancestor with the minimum ancestor-descendant rate change. Assuming that the ancestral rate is equal to 1, we have the relationship, (*r*_1_ × *r*_2_)^1/2^ = 1 [27]. We used the geometric mean, because relative rates could be very different from each other. We then project (recalibrate) *b*_1_ and *b*_2_ by determining the values of *r*_1_ and *r*_2_ which reconcile the two different estimates of *λ*_x_ (Fig. 1C).

The projected *b*_1_ is *b*_1_′ = *b*_1_ × (1/*r*_1_) and the projected *b*_2_ is *b*_1_′ = *b*_2_ × (1/*r*_2_). To determine the appropriate rate change factors, we first require that the root-to-X length (*λ*_X_) computed using *λ*_1_ and *b*_1_′, i.e., *λ*_1_ − *b*_1_′ = *λ*_1_ − *b*_1_ × (1/*r*_1_), and *λ*_X_ using *λ*_2_ and *b*_2_′, i.e., *λ*_2_ − *b*_2_ × (1/*r*_2_), be identical. Thus, we obtain the relationship, *λ*_1_ − *b*_1_ × (1/*r*_1_) = *λ*_2_ − *b*_2_ × (1/*r*_2_). Second, we use the constraint (*r*_1_ × *r*_2_)^1/2^ = 1, to solve for *r*_1_ = 0.93 and *r*_2_ = 1.08 in the current example. Similarly, for node Y, we calculate *r*_3_ and *r*_4_, which gives *r*_3_ = 0.99 and *r*_4_ = 1.01.

In the next step, we compute the relative rates of *b*_X_ and *b*_Y_, i.e., *r*_X_ and *r*_Y_, respectively. We similarly use projected branch lengths, *b*_*i*_′, and projected root-to-tip lengths, *λ*_*i*_. Here, we use the shortest root-to-tip length in each lineage of X and Y, because it is closest to a known sampling time from the root. Because *x*_1_ and *x*_3_ give the shortest length in the lineages X and Y, respectively, *λ*_XY_ on lineage X is given by *λ*_1_ − *b*_1_′ − *b*_X_′, and lineage Y gives *λ*_3_ − *b*_3_′ − *b*_Y_′ (Fig. 1D). Thus, we seek to enforce *λ*_1_ − *b*_1_′ − *b*_X_′ = *λ*_3_ − *b*_3_′ − *b*_Y_′. Given that (*r*_X_ × *r*_Y_)^1/2^ = 1, we can calculate *r*_X_ = 1.07 and *r*_Y_ = 0.93. Note that we previously assigned *r*_X_ equal to 1, as the ancestral rate of *b*_1_ and *b*_2_ correspond to *r*_X_. Similarly, *r*_Y_ was assigned to be 1. Therefore, the relative rates in the descendant branches are rescaled. For example, the new relative rate for the branch leading *x*_1_ becomes *r*_1_new_ = *r*_1_ × *r*_X_ = 0.93 × 1.07 = 1.00. Accordingly, projected branch lengths in the descendant lineages are rescaled, e.g., *b*_1_′ = *b*_1_ × (1/*r*_1_new_).

Since all tip branch lengths are now projected, we can obtain projected lengths from root to each internal node, i.e., *λ*_X_, *λ*_Y_, and *λ*_XY_. For example, *λ*_X_ is equal to be 1.66 [= *λ*_1_ − *b*_1_′ = *λ*_1_ − *b*_1_ × (1/*r*_1_new_) = 3.48 − 1.82 × (1/1.00)] (Fig. 1E). Using *λ*_X_, *λ*_Y_, *λ*_XY_, and the regression line, *λ* = 2.479 × *t*_*i*_ − 4957 (Fig. 1B), we obtain divergence times at the nodes XY, X, and Y to be 1999.9, 2000.3, and 2000.4, respectively (Fig. 1F).

### Performance evaluation using the simulated data sets

We evaluated the performance of RTDT by analyzing simulated data sets, as the true sequence divergence times are known for these data. We used the correct tree topology (branching pattern) in all our analyses because we wish to compare the true and estimated times, which is not possible for all the nodes in the true tree if the inferred tree contains errors. Also, we did not wish to confound the impact of errors in topology inference with that of the time estimates. In the same vein, we used the correct nucleotide substitution model to keep our focus on the accuracy of the time estimation methods, rather than on the problems encountered by the misspecified nucleotide substitution model.

In total, we analyzed 700 simulated viral phylogenies. In the following, however, we first present results from computer simulations conducted using parameters and tree topology derived from a DNA sequence alignment of subtype F HIV-1 [30] – a representative dataset with 154 strains with various sampling times (years 1987 - 2007; Fig. 2A) which was previously analyzed using BEAST. We generated two collections of simulated datasets using this model phylogeny. In one, evolutionary rates varied independently from branch to branch (IBR model) and in the other rates were autocorrelated between ancestor and descendant branches (ABR model). We also generated a collection of simulated datasets in which the expected evolutionary rates were the same for all branches (constant branch rates, CBR model), to serve as the baseline model. Fifty replicates were simulated with each clock model (CBR, ABR, and IBR).

**Figure 2.**
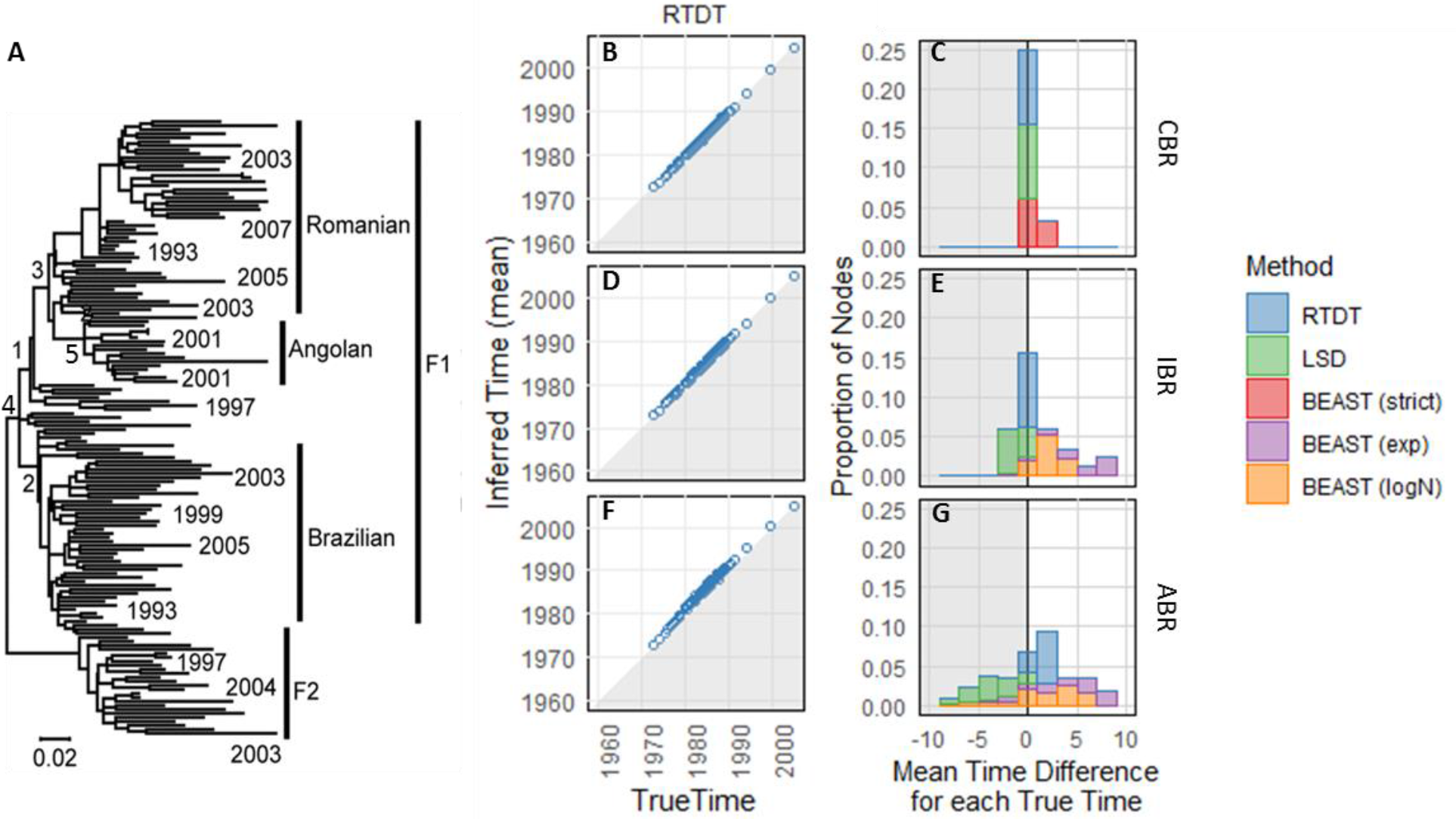
RTDT estimates (average node time) for computer simulated datasets. (A) Phylogeny of HIV-1 subtype F was used as the model tree. A few sampling times are shown at the tips. The number along a node is the node ID corresponding to nodes of importance in the original study [30]; see also Table 1. (B-G) Average node time estimates by RTDT for datasets simulated under (B) CBR clock model, (D) IBR clock model, and (F) ABR clock model. Stacked histograms showing average time difference from each correct time are given in panels C, E, and G for CBR, IBR, and ABR, respectively. These averages were means from 50 simulated datasets (replicates) at each node. For BEAST, we used a strict rate model for the analyses of datasets with CBR, and exponential (exp) and log-normal (logN) rate models were used for IBR and ABR data sets (C, E, and G). The shaded areas indicate that the average estimates are older than the actual times (B-G).

We used RTDT, BEAST, and LSD to compute timetrees with the correct nucleotide substitution model and the true topology. For each method, fifty time estimates were generated for each node in the model phylogeny. First, we examined the performance of RTDT, which are presented for CBR, ABR, and IBR models in Fig. 2B, 2D, and 2F, respectively. RTDT produced average time estimates that were very similar to the actual time for each node, i.e., RTDT performed well in estimating divergence times for this model tree. The percent deviation between the true time and the average estimated time (Δt) for all the nodes was close to zero (Fig. 2C, 2E, and 2G), suggesting that RTDT estimates are mostly unbiased.

We found LSD to also performed well for the CBR and OBR data sets, however, LSD was less accurate than RTDT for the ABR data sets (Fig. 2G and 3D). LSD estimates for ABR datasets yielded overly older dates, a problem that became more severe for deeper divergences. This is probably because LSD assumes rates to be independent among branches [16].

In BEAST analyses, we used strict clock model for the CBR data sets, so it showed an excellent performance for the CBR data sets. BEAST also performed well for IBR databases, but there was a small bias (Fig. 2 and 3) that may be attributed to the fact that BEAST assumed a log-normal distribution of branch rates but the simulations utilized a truncated uniform of rates. Such a bias became more extensive in the analysis of ABR datasets in which rates were autocorrelated (Fig. 2 and 3), because BEAST assumes branch rates to be not correlated. BEAST produced much older dates for deeper divergences, a pattern that was also seen in the LSD analyses, possibly because both methods assume independence of rates. The use of an exponential distribution of rates in BEAST performed worse for both IBR and ABR data sets (Fig. 2 and 3). Overall, RTDT outperformed BEAST and LSD on the ABR data sets, and showed a similar performance for IBR and CBR datasets.

**Figure 3.**
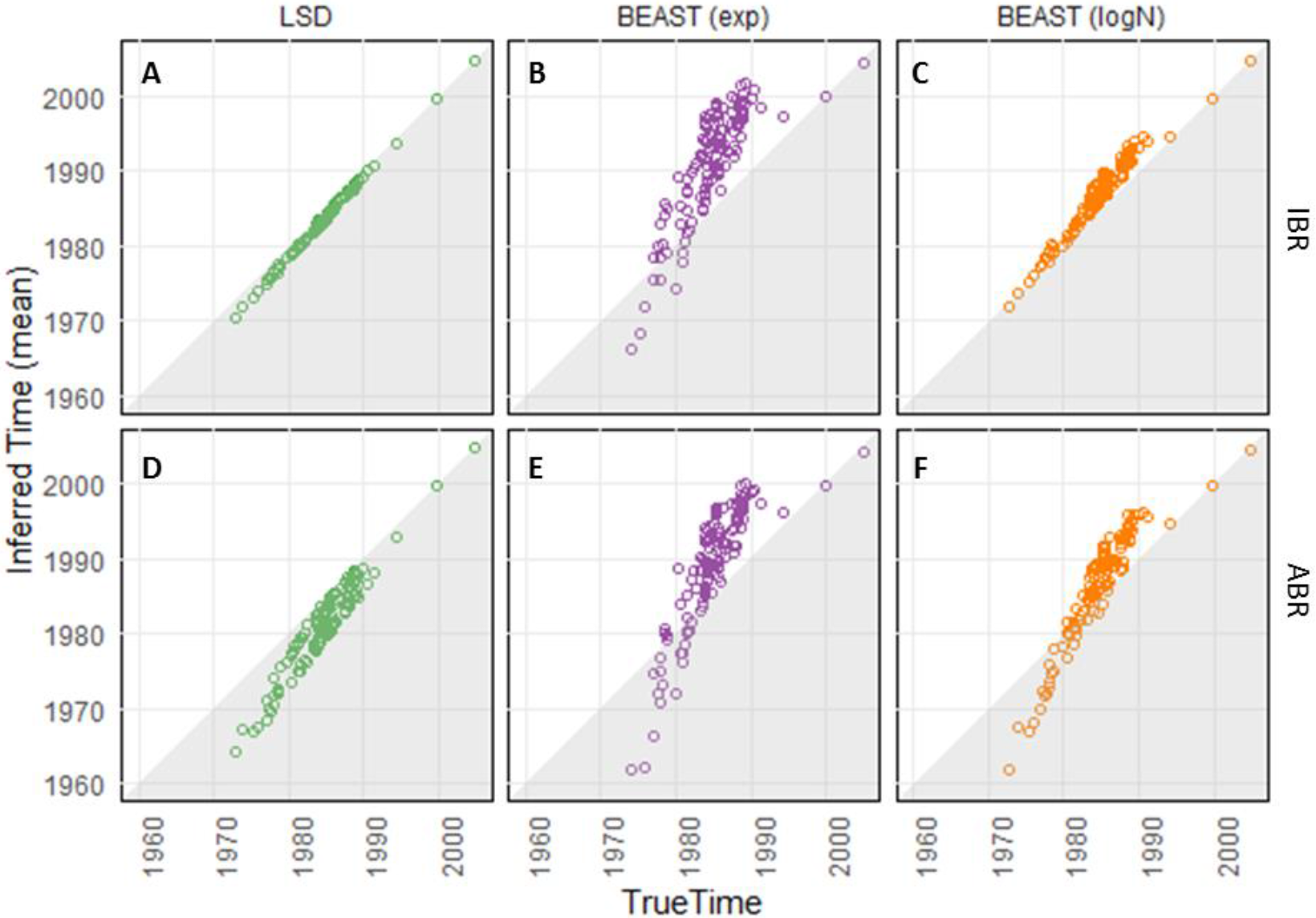
Average node time estimates of LSD and BEAST for datasets simulated following the model tree in Figure 2A. We generated datasets under IBR model (A-C) and ABR model (D-F). For BEAST, we used exponential (exp) and log-normal (logN) distributions of rates. The shaded areas indicate that the average estimates are older than the actual times. The results of RTDT are presented in Figure 2.

Although the average of node time estimates across replicates showed an excellent agreement with the correct node time for RTDT, the estimates varied extensively among replicates (Fig. 4). We found that standard deviations of estimated times were the smallest for recent divergences in all the methods, because they are the closest to the tips. As expected, the distribution of the oldest divergence times showed a much larger spread, because they were furthest from the tips in the model tree. These divergences span many branches that experienced extensive evolutionary rate changes over time. Consequently, accurate time estimation of deep divergences was generally difficult, especially when the branch rates were autocorrelated.

**Figure 4.**
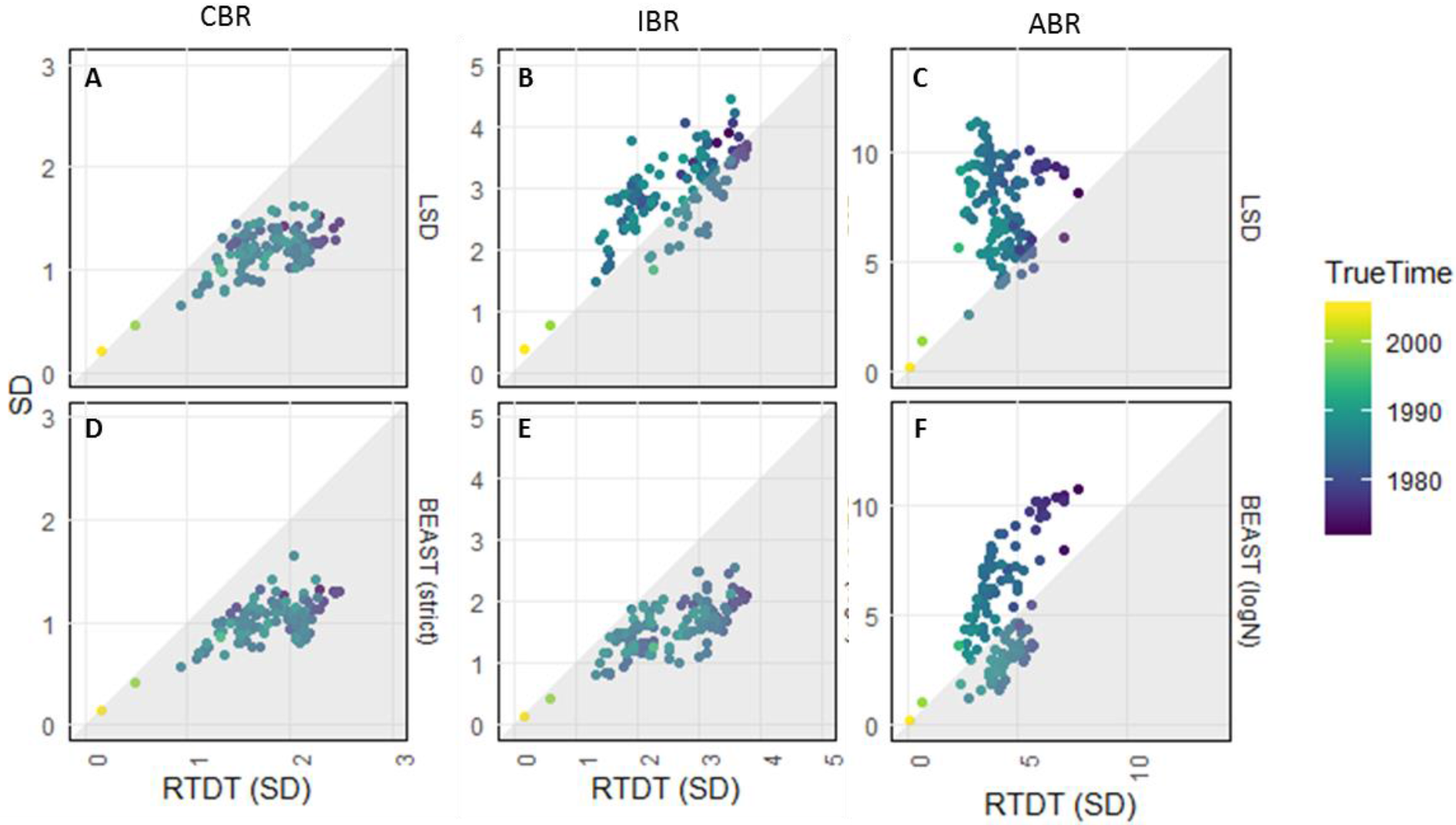
A comparison of standard deviations (SDs) of node times. (A-C) The comparison between RTDT and LSD on CBR (A), IBR (B), and ABR (C) datasets. (D-F) The comparison between RTDT and BEAST with strict clock rate model on CBR (A) and log-normal rate model on IBR (B) and ABR (C) datasets. Each point is an SD derived from a node with time estimates of 50 replicates. The color of a point indicates its true node times. Note that the average node time is presented in Figure 2 and 3.

Next, we tested the performance of timetree methods for datasets simulated using a larger (289 taxa) Influenza A virus phylogeny (Fig. 5A)[15]. This phylogeny is dramatically different from the HIV-1 phylogeny in figure 2A because the influenza A phylogeny is more ladder-like and is highly unbalanced. We simulated 50 datasets and analyzed them using the correct model and phylogeny in RTDT, LSD, and MCMCTree. We used MCMCTree instead of BEAST because it was employed in the source publication [15] and because BEAST (log-normal model) analyses for many of the datasets data sets failed to converge even after a long running time.

**Figure 5.**
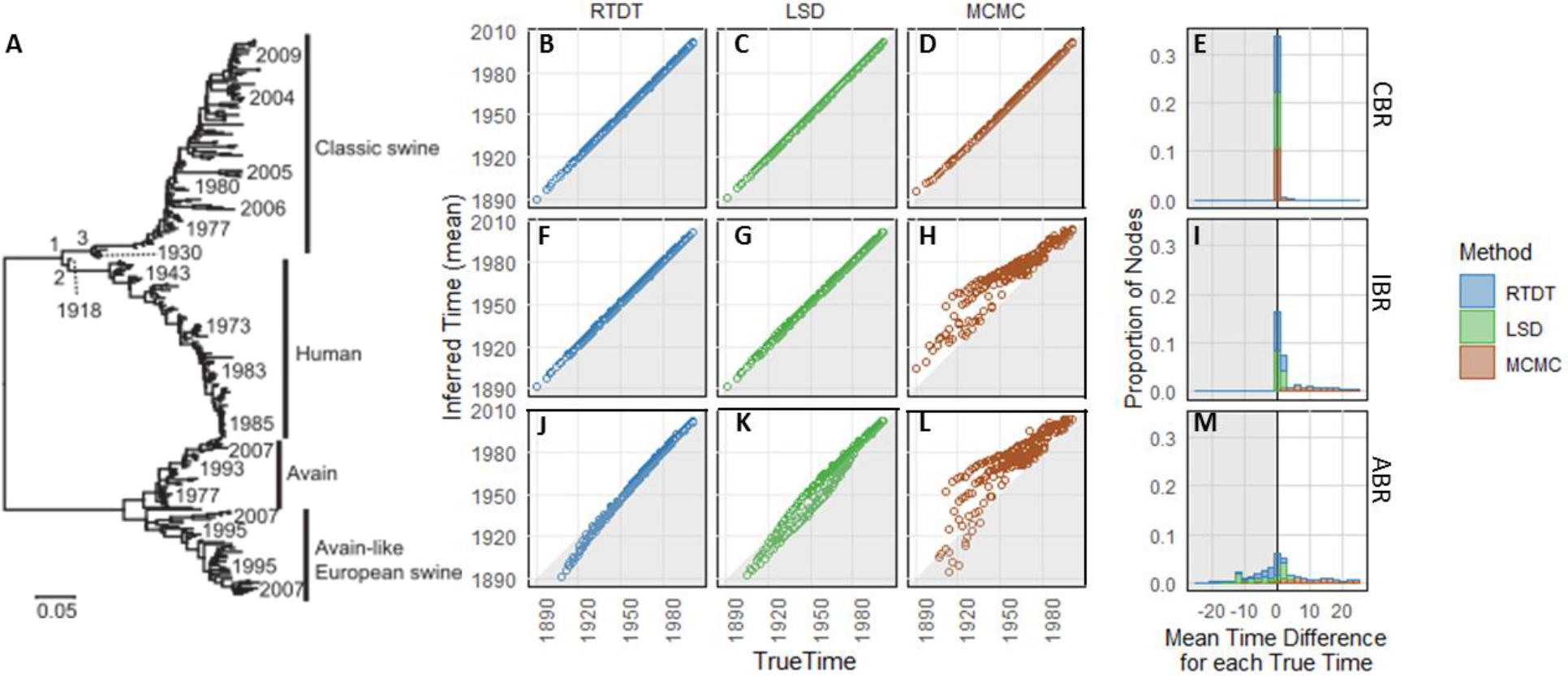
Comparison between RTDT, MCMCTree, and LSD. (A) Phylogeny of Influenza A. Sampling times are given for some tips. A number along a node is a node ID, which corresponds to those in Table 1. Fifty datasets were generated along this phylogeny with CBR, IBR or ABR. (B-M) Average node time estimates by RTDT, LSD, and MCMCTree (MCMC) for datasets with CBR (B-E), IBR (F-I), and ABR (J-M). Each time point is an average of 50 simulated datasets. MCMCTree was performed by using the correct branch rate model for each dataset. Average time difference from each true time is shown together in the form of stacked histograms (E, I, and M). The shaded areas indicate that the average estimates are older than true times.

RTDT performed well for Influenza A virus model phylogeny (Fig. 5B - 5M), but it showed a tendency to infer older ages for the oldest divergences under the ABR model (Fig. 5J and 5M). The performance of MCMCTree was worse than RTDT for both IBR and ABR datasets, even though the correct clock model was assumed in MCMCTree analyses (Fig. 5H and L). LSD and RTDT performed similarly for CBR and IBR datasets. However, for ABR datasets, LSD performed worse than RTDT for intermediate dates and better than RTDT for the deepest divergences (Fig. 5K). Therefore, RTDT and LSD were better than MCMCTree, but their performance was far from perfect. Overall, times estimated for the deepest nodes in ladder-like unbalanced trees must be interpreted with caution when branch rates are autocorrelated.

We next evaluated the performance of RTDT, LSD, and BEAST for datasets that mimic intra-host evolution (Fig. 6). We used To et al. [16] data, who simulated such intra-host datasets in which multiple strains are sampled at the same time. These strains may belong to the same clade (e.g., Fig. 6H) or different clades (e.g., Fig. 6A). Each dataset consisted of 110 sequences that were 1,000 bases long, and rates varied independently among branches [16]. Each simulated phylogeny was different from each other. In these datasets, many tips share the same sampling dates, and the number of different sampling dates is small (3 or 11 different dates).

**Figure 6.**
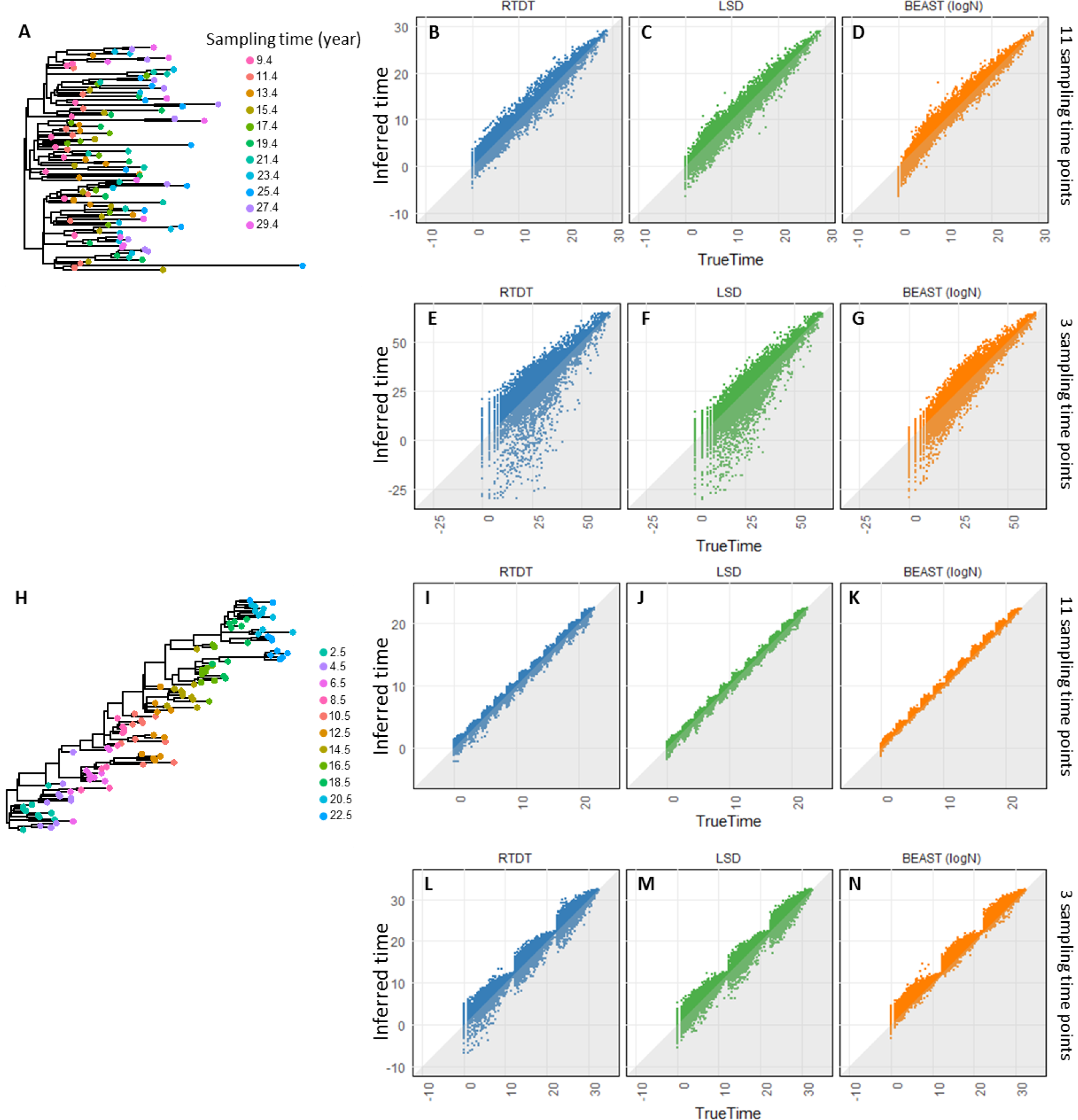
Comparison between RTDT, BEAST, and LSD using simulated datasets with a small number of sampling time points. (A) An example of HIV-like phylogeny. Tips are colored based on the sampling times. In this phylogeny, the root age was set to year of 0 (true age). Datasets were generated with independent rates. (B-G) Node time estimates by RTDT, LSD, and BEAST (log-normal rate model) for datasets with eleven sampling time points (B-D) and three sampling time points (E-G). (H-N) An example of Influenza-like phylogeny (H) and node time estimates (I-N). The shaded areas indicate that the average estimates are older than true times.

In the analysis of To et al.’s datasets with phylogenies similar in shape to the HIV-1 model tree (Fig. 6A; Fig. 2A), RTDT, LSD, and BEAST showed accuracies consistent with those observed for the HIV-1 model tree (Fig. 2 and 3) when the number of sampling time points was large, i.e., eleven time points (Figs. 6B - 6D). However, the situation became worse for all the methods, on data with only three sampling times (Fig. 6E-G), yielding much higher variances in node times estimates, especially for the deep nodes. Also, all methods inferred substantially earlier times for the deep nodes for a few datasets, which suggests loss of signal.

For ladder-like phylogenies in To et al.’s datasets (e.g., Fig. 6H), sequences were temporally clustered. Results from 11 sampling points show an excellent linear relationship with the true times (Fig. 6I-K). However, the relationship showed an undulating pattern of high and low dispersion, with the low dispersions observed for nodes that were located close to the tips. For these datasets, BEAST (log-normal rate model) frequently estimated divergences to be much younger, as compared to RTDT and LSD. With fewer sampling points, the pattern becomes clear because bias becomes higher (Fig. 6N). Overall, all methods showed limited accuracies on phylogenies in which the number of sampling dates was much smaller than the number of samples.

### Analyses of empirical data sets

We also explored some empirical datasets (Supplementary material Fig. S1 and Table 1) to examine how the patterns of published time estimates would have differed if RTDT was used instead of BEAST [14] or MCMCTree [31] programs. We begin with HIV-1 subtype F dataset because we used phylogeny and other evolutionary characteristics of this dataset as a model for our HIV simulation study (Fig. 2A). We found that estimates obtained by Mehta et al. [30], with BEAST using a log-normal rate model, were always older than those produced by using RTDT (Table 1). This result was consistent with our simulation results, as all of these nodes are located deep in the HIV-F phylogeny (Fig. 2A), for which BEAST is expected to show a tendency to infer older dates on ABR data (Fig. 3F). CorrTest [29] of this empirical dataset supported an autocorrelated clock model (*P* < 0.05). Therefore, one may prefer node ages produced by RTDT. Fortunately, RTDT dates do not contradict many of the biological scenarios presented by Mehta et al. [30], because the 95% highest posterior density (HPD) intervals of BEAST estimates generally included RTDT estimates (e.g., 1972-1983 and 1987, respectively for node 3).

**Table 1:** Empirical datasets used in this study

| Table 1: Empirical datasets used in this study |  |  |  |  |  |
| --- | --- | --- | --- | --- | --- |
| Virus | Node* | Time Estimates (year) |  | Clock model | Reference |
|  |  | RTDT | Bayesian | CorrTest |  |
| HIV-1 Subtype F (154 sequences, 1293 bps)<sup>a</sup> |  |  |  | Autocorrelated | Mehta, et al. (2011) |
|  | Node 1 | 1987 | 1975 |  |  |
|  | Node 2 | 1987 | 1980 |  |  |
|  | Node 3 | 1987 | 1978 |  |  |
|  | Node 4 | 1987 | 1973 |  |  |
|  | Node 5 | 1991 | 1984 |  |  |
| HIV-1 Subtype D (24 sequences, 2173 bps)<sup>a</sup> |  |  |  | Autocorrelated | Parczewski, et al. (2012) |
|  | Node 1 | 2003 | 2001 |  |  |
|  | Node 2 | 2000 | 1999 |  |  |
|  | Node 3 | 1995 | 1997 |  |  |
|  | Node 4 | 2006 | 2003 |  |  |
| HIV-1 Subtypes B/D (38 -133 sequence, 1497 - 8877 bps)<sup>a,d</sup> |  |  |  | Mixed<sup>x</sup> | Worobey, et al. (2016) |
|  | Node 1 | 1960 - 1969 | 1966 - 1969 |  |  |
|  | Node 2 | 1964 - 1971 | 1969 -1972 |  |  |
|  | Node 3 | 1966 - 1973 | 1969 - 1974 |  |  |
| HIV-2 (33 sequences, 1107 bps)<sup>b</sup> |  |  |  | Autocorrelated | Stadler and Yang (2013) |
|  | Node 1 | 1983 | 1938-1941 |  |  |
|  | Node 2 | 1985 | 1956 |  |  |
|  | Node 3 | 1985 | 1961-1964 |  |  |
| Rabies (67 sequences, 1350 bps)<sup>a</sup> |  |  |  | Independent | McElhinney, et al. (2011) |
|  | Node 1 | 1901 | 1885 |  |  |
|  | Node 2 | 1924 | 1917 |  |  |
|  | Node 3 | 1937 | 1931 |  |  |
|  | Node 4 | 1945 | 1941 |  |  |
| Influenza A (289 sequences, 1710 bps)<sup>c</sup> |  |  |  | Autocorrelated | Stadler and Yang (2013) |
|  | Node 1 | 1912 | 1813-1910 |  |  |
|  | Node 2 | 1915 | 1832-1914 |  |  |
|  | Node 3 | 1928 | 1889-1926 |  |  |
| a: BEAST with lognormal rates |  |  |  |  |  |
| b: MCMCtree with constant and autocorrelated clock models |  |  |  |  |  |
| c: BEAST with lognormal rates and MCMCtree with constant, independent, and autocorrelated clock models. The range of estimated times based on these different methods was given. |  |  |  |  |  |
| d: The range of time estimates was obtained based on eight different subdatasets. |  |  |  |  |  |
| x: Five datasets showed autocorrelated rates and three independent rates. |  |  |  |  |  |
| *: Node IDs were given in **Figures. 2, 5, and 7.** |  |  |  |  |  |

We next examine results for Influenza A viral dataset, which served as a model for our influenza simulations (Fig 5A). Different Bayesian methods produced different time estimates, and an autocorrelated rate model in MCMCTree always produced much older times than the other rate models in MCMTree and BEAST (log-normal rate model). RTDT estimates were younger than all the Bayesian estimates (Table 1), but the difference was small when considering BEAST with log-normal rate model, e.g., 1813, 1898, and 1912 by MCMCTree with the autocorrelated model BEAST (log-normal rate model) and RTDT, respectively for node 1. This result was also consistent with our simulation results. An ABR clock model fits this data set according to CorrTest (P<0.001), and our simulations already showed that MCMCTree with an autocorrelated model has a stronger tendency to generate older dates for deep nodes (Fig. 5L) as compared to RTDT (Fig. 5F).

Results from the analysis of two other HIV-1 datasets – *subtypes B/D [32] and subtype D [33]* – showed high concordance between RTDT and Bayesian analyses (Table 1). For Rabies data, although BEAST estimates were slightly older than RTDT, we found that these RTDT estimates were within the 95% HPD intervals. The only exception was HIV-2, in which RTDT produced node times that were much younger than those from MCMCTree analysis. This discrepancy occurred because this data did not contain much temporal structure, as the root-to-tip lengths and sampling times did not show a good positive correlation (Supplementary material Figure S2). Tip-dating methods are known to be adversely affected by such data and their use is generally not recommended [34, 35].

Overall, RTDT may be preferred in empirical data analysis. This choice is made easier by the fact that RTDT is orders of magnitude faster than the Bayesian methods. For example, the Influenza A virus dataset with 289 sequences was analyzed in only a few minutes by RTDT, but it took BEAST 4.4 days when using a lognormal distribution of rates.

## DISCUSSION

We have presented a new relaxed-clock method to estimate times of sequence divergence using temporally sampled pathogenic strains. The new method (RTDT) is based on the relative rate framework in the RelTime method [27] but represents a significant advance of this framework as it removes the requirement that the sequences sampled to be contemporaneous. In RTDT, there is no need to specify autocorrelation vs. independence of rates or to select a statistical distribution for rates, which is an advantage over Bayesian methods where such information is required a priori.

In the analysis of computer simulated data, RTDT performed similar or better than the Bayesian approaches tested, while Bayesian methods are the most widely used methods in empirical data analyses [34]. We found that Bayesian methods produced much older time estimates for the deepest nodes than RTDT when the evolutionary rates were autocorrelated. The worse performance of BEAST on ABR data can be attributed to the clock model violation because BEAST assumes that rates vary independently among branches. This result is consistent with Wertheim et al. [25], who reported that Bayesian methods produced erroneous node times when evolutionary rates are lineage (clade) specific, similar to what was used for our ABR simulations.

Also, we found another non-Bayesian method (LSD) to perform worse than RTDT for datasets with autocorrelation of rates (Fig. 3 and 5), likely because LSD assumes that the rate variation among branches in the phylogeny follows a normal distribution, which may not be satisfied because log-normal distribution may fit the data better when the branch rates are autocorrelated. Nevertheless, LSD performed similar or better than the Bayesian approaches, a pattern that has been seen in the past as well [16].

As mentioned earlier, we assumed the correct phylogeny as well as the correct substitution pattern in our computer simulations. However, clearly, inferred phylogenies contain estimation errors and the nucleotide substitution pattern selected may be suboptimal, both of which will impact the accuracy of time estimates. A comprehensive investigation is necessary to better evaluate the robustness of RTDT, BEAST, and LSD in those situations, which is beyond the scope of the current article. However, it is interesting to note that in the analysis of HIV-1 subtype B/D datasets, we observed similar divergence times for these datasets (Table 1), which suggests that topological errors within strains did not have a large adverse impact. Nevertheless, robust inference of evolutionary relationship of strains or sequences of interest may not be possible under certain situations [36, 37], and in such cases, the estimation of divergence times will likely be misleading. Similarly, unreliable branch length estimates will result in poor time estimates, which has been previously highlighted in ref. [24]. In conclusion, RTDT can produce similar or better results than other methods, including Bayesian and non-Bayesian approaches. RTDT method is implemented in the cross-platform MEGA X software that is freely available from http://www.megasoftware.net.

## MATERIAL AND METHODS

### Collection and Analyses of Empirical Datasets

Nucleotide sequence alignments and sampling time information of nine different viruses (see Table 1 for the detail) were obtained from the supplementary information [15], Dryad Digital Repository (https://datadryad.org/) [32], or the authors [30, 33, 38]. Note that the HIV-1 Subtype B/D data [32] was composed of eight datasets, in which each dataset contained sequences of genes (env, gag, or pol) or the full genome with various numbers of sequences.

### Computer Simulation

We simulated nucleotide sequence alignments along viral timetrees obtained from the original studies (subtype F HIV-1 [30] and Influenza A [15]) and the respective nucleotide substitution rates, transition/transversion ratio, CG contents, sequence lengths, and substitution models. The nucleotide substitution rates were obtained from these original studies (3.2 × 10^−3^ and 1.7 × 10^−3^ per site per year for subtype F HIV-1 and Influenza A, respectively). The average transition/transversion ratios were 2.7 and 2.6, respectively, and the average CG contents were 38% and 41%, respectively. The nucleotide sequence lengths simulated were the same as in the original datasets (1,293 bps and 1,710 bps, respectively). Note that the tips of branches on the timetrees were truncated according to the sampling times, which were also obtained from the original studies.

Using the Seq-Gen software [39] under HKY substitution model [40], 50 alignments were generated for each timetree with the constant rate (CBR), randomly varying rate (IBR), and autocorrelated rate (ABR) among branches, following the methods in Tamura et al. [28]. For IBR, each mutation rate was drawn from a uniform distribution with the interval ranging from 0.5*r* to 1.5*r*, where *r* is the original mutation rate in the simulation above. For ABR, the rate variation was autocorrelated between ancestral and descendant lineages. The rate of a descendant branch was drawn from a lognormal distribution with the mean rate of the ancestral branch and the variance equal to the time duration, in which the autocorrelation parameter, *v* in Kishino et al. [41], was set to 1. Among these datasets, we removed datasets when it included identical sequences between different taxa, because all sequences should be distinct in actual empirical data. In total, we used 50, 49, and 43 datasets for Subtype F HIV-1 with CBR, IBR, and ABR, respectively, and 50, 50, and 38 datasets for Influenza A virus with CBR, IBR, and ABR, respectively.

We obtained 400 LSD datasets (IBR) from the LSD website [http://www.atgc-montpellier.fr/LSD/], which excluded 77 datasets because they contained at least two identical sequences.

### Analyses of simulated data

For each simulated alignment, each node time was estimated using the correct tree topologies and sampling times, which were obtained from the original studies. RTDT estimates were obtained using MEGA-CC [42] with the HKY nucleotide substitution model [40] with gamma-distributed rate heterogeneity among sites [43] because this option is widely used.

The same substitution model was used in the Bayesian methods. In BEAST [v1.8.0; 14], the strict clock model was used for analyzing CBR datasets, and independent (lognormal and exponential) branch rate model was used for analyzing IBR and ABR datasets. The constant population size model was selected for the coalescent tree prior. The number of steps that MCMC made was 10,000,000 steps, and trees were sampled every 1,000 steps. To evaluate if large enough genealogies (trees) were sampled, we used the TRACER software [44] and confirmed that the number of independent information in the sampled posterior values (effective sample size; ESS) was at least 200. Since analyses using log-normal and exponential distributions of rate did not show at least 200 ESS, we used 100,000,000 MCMC steps for all the datasets. Among sampled trees, we excluded the first 10% of the trees as burn-in and computed the mean height of each node along the true tree topology using the TreeAnnotator software, which is implemented in the BEAST software.

Datasets generated along influenza A data were analyzed by using MCMCTree [PAML4.7; 31] because the source publication used MCMCTree. The default parameters were used, i.e., root age prior was between 50 and 200 years ago with the violation probabilities of 1%, and the time prior for the nodes in the tree was constructed using birth-death process. Discarding the first 20,000 iterations, 200,000 iterations were made, and trees were sampled every two iterations. Strict, independent, and autocorrelated clock model were used for analyzing datasets generated with the CBR, IBR, and ABR, respectively.

To obtain LSD [16] estimates, we first estimated branch lengths using the Maximum Likelihood method with HKY nucleotide substitution model under MEGA-CC [42], because LSD required a phylogeny with branch lengths as input. Along the phylogeny and sampling time information, each node time was inferred using the temporal constraints for node time estimates and considering the variance of branch length estimates, with the default parameters (lower bound for the rate is 0.00001 and parameter of variances is 10).

## Acknowledgments

We thank Qiqing Tao for critical comments. This work is supported by grants from NIH (R01GM126567-01), National Science Foundation (ABI 1661218), National Aeronautics and Space Administration (NASA, NNX16AJ30G), Pennsylvania Department of Health (TU-420721), and Tokyo Metropolitan University (DB105).

## Supporting information

**S1 Figure. Phylogenies from the published literature for empirical datasets.** Branch lengths were the number of substitutions. Sampling times were indicated for a few sequences. A number along a node is a node ID, which corresponds to that in Table 1. Those node times were reported in the original study.

**S2 Figure. Root-to-tip branch length and sampling time for HIV-2 data.** The trend line is *y* = 0.0044*x* − 8.5 (*R*^2^ = 0.20).

---

**Figure S1.**
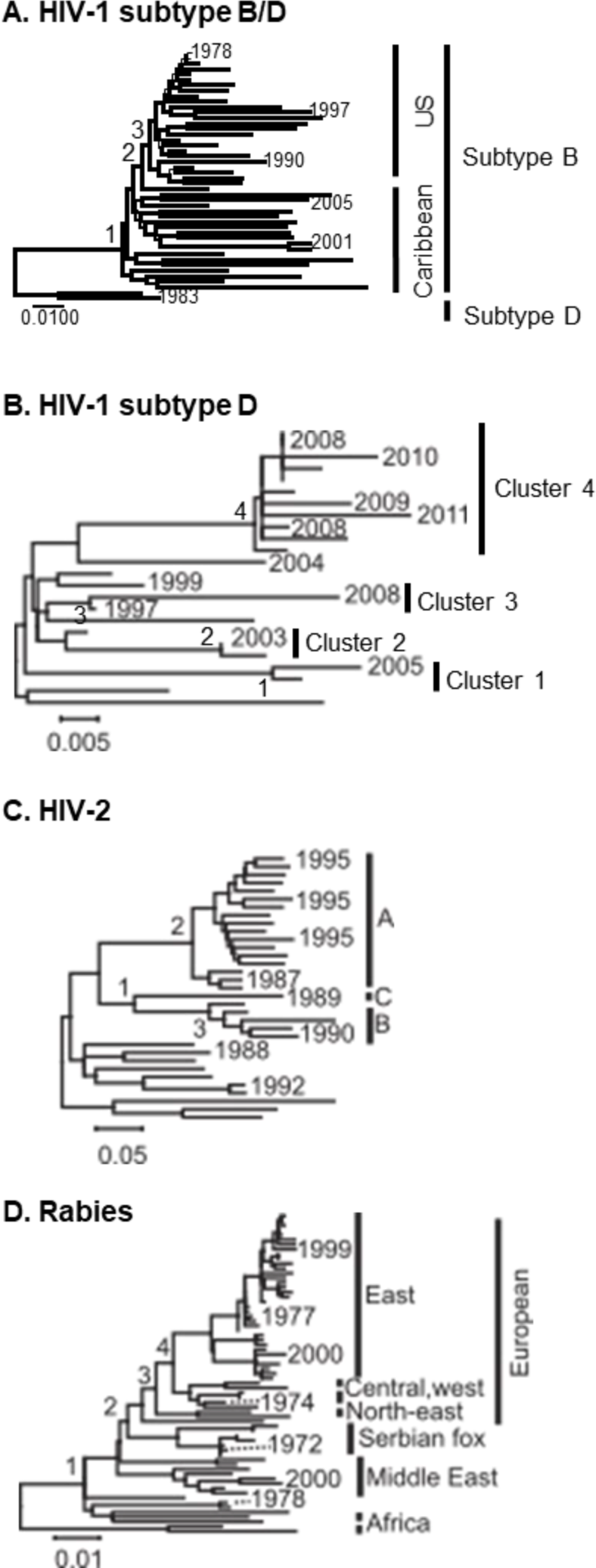
Phylogenies from the published literature for empirical datasets. Branch lengths were the number of substitutions. Sampling times were indicated for a few sequences. Numbers are node IDs, and those node times were reported in the original study (Table 1).

**Figure S2.**
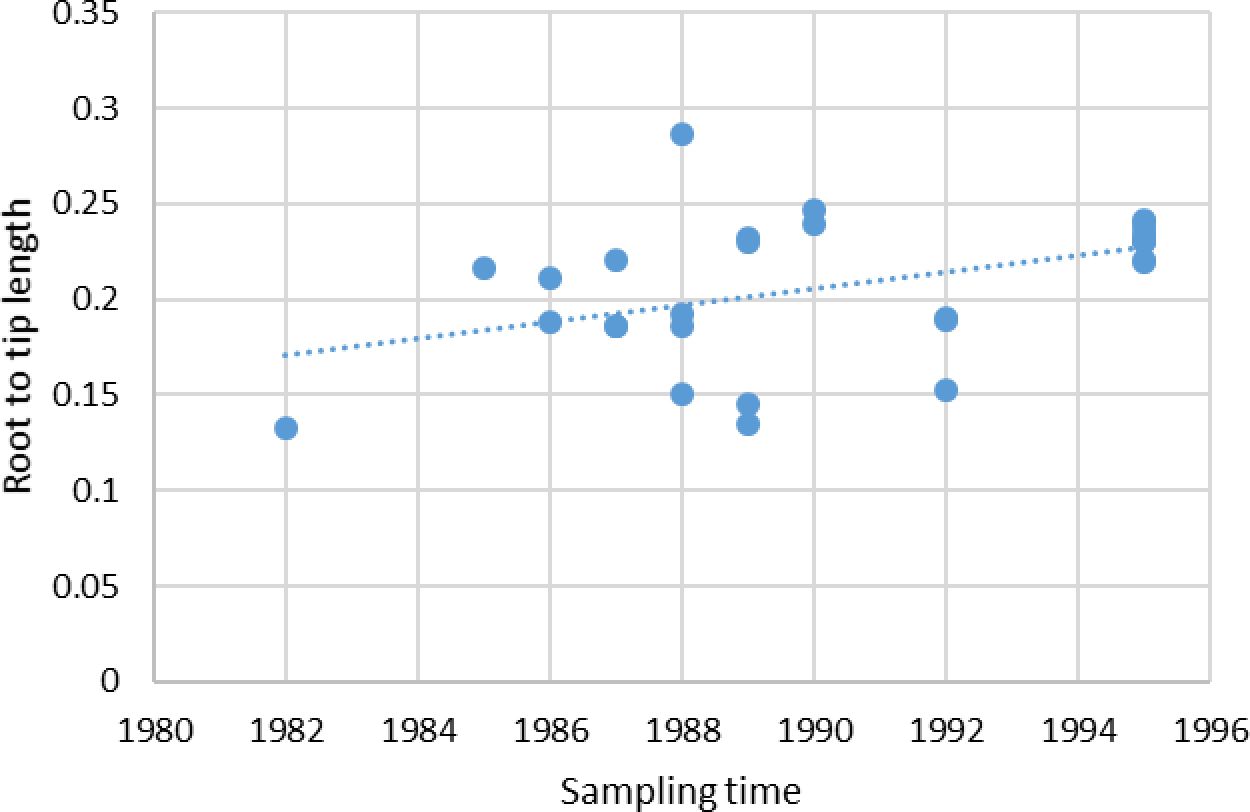
Root-to-tip branch length and sampling time for HIV-2 data. The trend line is *y* = 0.0044*x* − 8.5 (*R*^2^ = 0.20).

